# Quantifying per-match Reliability in Library Matching for Untargeted Metabolomics Workflows

**DOI:** 10.64898/2026.07.30.741704

**Authors:** Esteban Charria-Girón, Jorick van IJcken, Larissa Della Vedova, Laura Rosina Torres-Ortega, Justin J.J. van der Hooft

## Abstract

Tandem mass spectrometry has become central to untargeted metabolomics. The translation of unknown spectra into biological insight depends on assigning chemical identities to detected metabolites. Structural characterization typically begins with mass spectral library matching, in which experimental spectra are compared against reference libraries and candidate annotations are ranked by their spectral similarity to the query. As spectral libraries and experimental datasets grow, however, more candidates achieve comparable similarity scores for a single query, and similarity scores give no indication of how reproducible a candidate match is or how sensitive it is to the underlying fragment evidence. Existing false-discovery-rate approaches can indicate annotation error at the dataset level but do not provide a per-match estimate of reliability. Here, we introduce a SpecReBoot-inspired query-focused bootstrapping approach that resamples the fragment evidence of each query spectrum. This approach relies on recomputing query similarity to candidate library spectra across bootstrap replicates, which provides a statistical distribution of scores rather than a single value. From this distribution we define the match support, a per-match reliability estimate quantifying the reproducibility of a match under spectral perturbation, together with measures of ranking stability that describe how often a candidate remains among the top-ranked matches across replicates. Applied to a forensic drug-of-abuse case, match support distinguished previously identified annotations from high-scoring false positives: a distinction cosine similarity failed to make. Furthermore, match support values remained stable as the reference library was expanded, whereas ranking stability metrics shifted significantly. In a cross-instrument endogenous metabolite library search, match support further revealed metric-specific annotation behavior, identifying metabolites consistently supported across different similarity metrics, while flagging annotations whose reliability depended strongly on the chosen scoring metric. Benchmarking against a natural-product reference library demonstrated that ranking based on match support values promoted true matches by four ranks on average compared with cosine-based ranking, without promoting analogs. Under controlled spectral perturbation experiments, match support flagged incorrect annotations with an AUROC of 0.75, whereas the cosine similarity score alone of the same match reached only 0.56. Query-focused bootstrapping thus provides a practical, per-match measure of annotation reliability, bringing the field a step toward reliable annotations at scale. We anticipate that incorporation of our annotation reliability scoring into computational metabolomics workflows will further promote the growth of spectral libraries and enhance their applicability across scientific disciplines.

## INTRODUCTION

Tandem mass spectrometry (MS/MS) has become a key technology for the routine analysis of biological and chemical samples, bringing insight into the chemical diversity of complex mixtures and transforming the way we study the chemistry of life.^1,2^ Yet, the success of untargeted metabolomics as a central approach for studying complex samples and facilitating biological discovery depends on our ability to assign identities to the large number of MS/MS spectra acquired without prior knowledge of their chemical composition.^3,4^ In most untargeted metabolomics workflows, the first step toward establishing the identity of detected ions or features is to match experimental spectra against public or in-house spectral libraries, in what is referred to as library matching.^5,6^ The resulting spectral matches are ranked by their spectral similarity to the query spectrum, and these annotations are either accepted or rejected based on heuristics such as the similarity threshold or the minimum number of matching peaks.^6^ Nevertheless, a more fundamental limitation persists: spectral matches do not indicate how well each hit is supported by the underlying spectral evidence.

Although strategies for estimating false discovery rates (FDRs) in library matching have been introduced, these methods have not yet been widely adopted by the community.^7−9^ FDR-based approaches generally provide a global error estimate by estimating the expected rate of incorrect annotations across a particular dataset, or across a selected set of accepted matches, rather than assigning a reliability value to each individual query–candidate pair. This distinction is important because a dataset-level FDR can indicate that the overall annotation list is controlled at a given error rate, while still leaving uncertainty about which specific annotations are robust and which are fragile. Target-decoy approaches, in which spectra are searched against a set of deliberately incorrect reference spectra so that the rate of chance matches can be estimated, have become the gold standard in other data-intensive fields such as proteomics;^10,11^ however, they have been of limited benefit in metabolomics because generating realistic decoy spectral libraries is challenging. Still, substantial progress has been made in adapting target-decoy approaches to tandem mass spectral data.^8,12^ At the same time, untargeted mass spectrometry profiles and public mass spectral libraries have grown rapidly in size (around a 60-fold increase in the last 10 years) and coverage in recent years, advancing our ability to study small molecules.^7^ As the number of candidate spectra increases, so does the chance of having more competitors with comparable similarity scores for a single query, raising the number of plausible but incorrect annotations. Realistic alternatives for evaluating the reliability of spectral library matches are therefore needed, as untargeted metabolomics is scaling towards increasingly large datasets and reference libraries.

We recently introduced the concept of confidence-aware mass spectral networking with SpecReBoot that systematically resamples fragment-level features to derive an empirical similarity distribution for each spectral pair, from which a bootstrap support value is assigned to each edge of the resulting molecular network.^13^ Based on the edge support, the network topology can be pruned of spurious spectral relationships, improving the chemical coherence of the resulting connected components. Unlike molecular networking, spectral library matching is a one-versus-all problem rather than all-versus-all. We hypothesize that adapting this bootstrapping-based framework to library matching will provide the per-match resolution that FDR estimation cannot. Hence, here we present a query-focused bootstrapping approach that introduces match support as a per-match measure of reliability, defined as the reproducibility of an annotation under bootstrap perturbation of the query’s fragment evidence, complemented by ranking-based stability measures defined within each query’s candidate set. By resampling the query’s fragment features, each match is associated with a statistical distribution of similarity scores rather than a single value, which aids in distinguishing false from true positives in data derived from complex biological samples.

We hypothesize that query-focused bootstrapping provides a per-match reliability signal that (i) distinguishes validated true from false positive annotations independently of the raw similarity score, (ii) remains stable as the reference library grows, whereas ranking stability approaches depend on the candidate space, and (iii) generalizes across similarity metrics, such that agreement among metrics identifies the most reliable annotations. To test these hypotheses, we first apply the approach to a forensic urine sample with known annotated drugs of abuse and their metabolites, providing a controlled scenario with previously supported annotations.^14^ Then, we extend the analysis to endogenous metabolites using multiple different spectral similarity metrics. Furthermore, we benchmark match support based reranking on publicly available natural product libraries of increasing size, where independent structural ground truth allows performance to be evaluated at scale rather than on isolated cases. Finally, to understand mechanistically when the approach succeeds and fails, we perform controlled spectral perturbation experiments that systematically degrade library spectra and track the response of match support relative to the raw similarity score.

## MATERIALS AND METHODS

### Datasets and accessions

#### Urine sample

The urine MS/MS data reprocessed in the present study were obtained from a previously published forensic toxicology investigation of a suspected driving-under-the-influence-of-drugs (DUID) case.^14^ The dataset consisted of liquid chromatography coupled to high-resolution mass spectrometry (LC–HRMS) spectra acquired from a urine sample collected during routine forensic casework and retrospectively analyzed in fully anonymized form. No additional sample collection, preparation, or instrumental acquisition was performed for the present study.

Raw LC–HRMS files were reprocessed using the mzmine 4 workflow.^15^ After feature detection and MS/MS spectral processing, the resulting tandem mass spectra were exported in Mascot Generic Format (MGF) and analyzed using the spectral library matching workflow described in the next sections. The compound annotations reported in the original study were used as reference identifications to assess the performance of our proposed strategy.^14^

#### Forensic spectral libraries

The first spectral library used in this study for the investigation of the DUID case was downloaded in MGF format from the public GNPS website (https://external.gnps2.org/gnpslibrary), specifically from the “Drugs of Abuse” section, and contained 480 MS/MS spectra. Subsequently, the GNPS-derived library and the MCE spectral library were downloaded from the same repository. The latter refers to a spectral library generated from MedChemExpress compound collections, including bioactive molecules, chemical scaffolds, and FDA-approved drugs. These three libraries were then merged into a single MGF file named “library_ABUSE_GNPS_MCE.mgf”, containing a total of 38,898 spectra. This library was then cleaned by matchms to obtain a cleaned library of 31,742 spectra “library_ABUSE_GNPS_MCE_cleaned_harmonized.mgf”. Both spectral datasets are also available at [https://zenodo.org/records/21507928].

#### Benchmark datasets

The NIH Natural Products Library dataset comprising 1,267 reference spectra (GNPS-NIH-NATURALPRODUCTSLIBRARY.mgf) was acquired from the public GNPS website (https://external.gnps2.org/gnpslibrary), while the MSn-COCONUT dataset containing 15,947 spectra was taken from the original SpecReBoot study.^13^ The respective datasets are available at [https://zenodo.org/records/21507928].

#### Endogenous Metabolite Spectral Library

To explore the generalizability of our library matching beyond forensic compounds, the same sample was additionally searched against an in-house spectral library of metabolically relevant small molecules detectable in human urine, enabling a complementary assessment of annotation stability across both drug-related and endogenous metabolite classes within a single biological sample. This library (101 reference compounds, 2,324 MS/MS spectra; 1,572 positive- and 752 negative-mode) was generated from authentic standards analyzed by pHILIC LC-MS/MS on a Q Exactive HF Orbitrap (Thermo Fisher Scientific) with electrospray ionization in both polarities and HCD fragmentation. Data-dependent acquisition (Top 5) followed the method described by Zaal et al.,^16^ with the following modifications for MS/MS events: resolution 30,000; AGC target 5 × 10⁵; maximum injection time 120 ms; isolation window 1.0 *m/z*; normalized collision energies 20, 30, and 40. Raw data were converted with MSConvert and processed using the mzmine 4 library generation workflow;^17^ the obtained spectral library was further cleaned and filtered as described in the accompanying processing notebook available at [https://zenodo.org/records/21394919]. The urine sample was searched against this library under cosine, modified cosine, Spec2Vec and MS2DeepScore similarities (*B* = 100, top-N = 100, score threshold 0.7, precursor tolerance 0.02 Da, analog search enabled).

### Implementation, required inputs and outputs

The query-focused library matching algorithm is embedded within the SpecReBoot ecosystem and operates on standard, GNPS-compatible MS/MS data. It starts with a query spectrum and a set of library spectra (typically MGF files) and returns a ranked table of candidate annotations enriched with bootstrap-derived metrics for each query.

#### Required inputs

1. 1.Query and library MS/MS spectra. Each spectrum should carry a stable, unique identifier in its metadata. Identifiers are resolved in priority order, spectrum_id, then feature_id, id, or scans, so that GNPS library entries are tracked by their unique spectrum_id (e.g. CCMSLIB00000079350) rather than the feature_id field, which is frequently exported as 0 for every spectrum in a GNPS MGF and is therefore not unique. Spectra lacking any usable identifier receive a generated fallback.
2. 2.A precursor *m/z* for each spectrum is recommended. When the query carries a precursor_mz, the library search can be first restricted to candidates within a selected precursor mass tolerance window. When it does not, the search proceeds against the full library, thus effectively becoming an analogue search; however, we note that similarity metrics such as modified cosine and Spec2Vec require a precursor mass for each spectral query.

#### Parameter selection and recommended settings

Default values were chosen to balance stability of the resulting metrics against computational cost and can be adjusted to the characteristics of a given dataset.

1. 1.The number of bootstrap replicates (*B*, default 100) determines the resolution of the resulting match support values, as this measure is quantized in steps of 1/*B*. When *B* = 100, the given resolution is 0.01, which is sufficient to distinguish candidates near the decision threshold, and in the DUID case varying *B* between 30 and 100 did not change the top-supported annotations. Lower values (*B* = 50) are suggested as a first screening where runtime dominates, as used in our perturbation experiments, at higher values a better resolution is expected at the cost of runtime.
2. 2.Only the top_n (default 100) candidates from the initial scoring are passed to the bootstrap phase, so this parameter sets the competitive field within which rank-stability measures are defined. Larger values give a more realistic picture of competition in large libraries but increase runtime proportionally. Smaller values (top_n = 10 to 20) are appropriate when only the highest-ranking candidates are of interest, for large perturbation grids, or if the library space is chemically sparse. However, whereas the actual inclusion of correct and false positive matches may depend on it, we note that the match support value is unaffected by top_n, since it is a property of the individual query–candidate pair, whereas rank-stability measures depend directly on it.
3. 3.Two decimal places for the binning resolution (decimals, default 2) correspond to 0.01 Da bins, which is appropriate for high-resolution data (QTOF, Orbitrap). Lower-resolution instruments require coarser binning, since bins narrower than the instrument’s mass accuracy will split fragments that should be treated as identical.
4. 4.Match support is defined relative to the score threshold (default 0.7), so its choice sets what counts as a supported replicate. We used 0.7 throughout, consistent with commonly applied (loose) cosine cutoffs in library matching. Because rank-stability measures are threshold- free, the two families of metrics can be interpreted together when threshold choice is uncertain.
5. 5.Precursor *m/z* tolerance (default 0.02 Da) mirrors the parent-mass tolerance in GNPS library matching. Recommended values are 0.01–0.05 Da for high-resolution calibrated data, 0.3–2.0 Da for low-resolution data, and 2.0 Da (the GNPS default) for mixed-instrument libraries.
6. 6.Analog search (default on) applies when fewer than top_n candidates fall within the precursor tolerance window, then remaining slots are filled with the highest-scoring spectra outside that window, flagged as analogs. This is useful for exploratory annotation of compounds absent from the library but should be disabled when only exact matches are of interest, since analogs share partial fragment evidence and therefore receive systematically lower support (see Results). Note also that enabling analog search increases runtime, since the candidate space is filled to top_n in every case.
7. 7.Any matchms compatible score metric is supplied directly, i.e., CosineGreedy, ModifiedCosine, FlashSimilarity, Spec2Vec, or MS2DeepScore, with no wrapping required. No default is provided, since the appropriate metric depends on the data and the annotation goal, fragment-based metrics (cosine, modified cosine) for matched-instrument searches, and embedding-based metrics (Spec2Vec, MS2DeepScore) where fragmentation may differ between query and library. As match support is defined relative to the chosen metric, support values are not directly comparable across metrics (see Results).

#### Expected outputs

For each query, a result object containing a per-candidate statistics table and a bootstrap top-hit frequency table. Per candidate, the table reports the original similarity score and rank, match support (fraction of replicates in which the candidate’s score meets score_threshold), the bootstrap score mean and standard deviation, top-1, top-3, and top-5 rank stability (fraction of replicates in which the candidate held that rank), and mean bootstrap rank, which are defined below. These are saved as CSV files, a top-hits table per query (all_top_hits.csv) enriched with library metadata read from the matched spectra (name, SMILES, InChIKey, formula, adduct, instrument type, collision energy), and a per-candidate statistics table (all_candidate_stats.csv).

### MS/MS preprocessing

All spectral data are first imported as MGF files and then harmonized and cleaned using the default matchms pipelines (DEFAULT_FILTERS and CLEAN_PEAKS) available and user- adaptable in the SpecReBoot codebase.^18,19^ This step normalizes intensities to unit height and applies the standard matchms cleaning steps: removal of peaks within ±17 Da of the precursor (excluding the precursor itself), retention only of peaks within ±50 Da of one of the six most intense peaks, and a cap of 1,000 fragment peaks per spectrum. These defaults settings can be adjusted to specific dataset characteristics and metadata completeness.

### SpecReBoot query-focused library matching algorithm

#### Query-focused bootstrap resampling

Our library matching approach builds on the resampling framework previously introduced for confidence-aware mass spectral networking,^13^ in which each spectrum’s fragment peaks are first discretized onto a shared *m/z* grid by binning, and the set of occupied bins across the spectra defines a global bin list *G*. In the networking scenario, *G* contains all spectra in an all-versus-all comparison. For library matching, we instead adopt a query-focused alternative in which the global bin list *G* is constructed from the query spectrum alone, so that each replicate resamples over bins informative for the query rather than over the combined *m/z* space of all candidates, which can be dominated by library-only peaks (Figure 1a). For each bootstrap replicate b, P bin indices are sampled uniformly with replacement from 0 to P, generating a unique sampled bin subset *G*⁽ᵇ⁾ ⊆ *G*, where each bin value represents a collection of binned original fragments. Because sampling is with replacement, on average 1 − (1 − 1/P)^P ≈ 63.2% of the global bins are retained in each replicate and the remaining ∼36.8% are dropped. The query spectrum and each candidate are converted into pseudo-spectra by retaining only those peaks whose discretized *m/z* value falls in *G*⁽ᵇ⁾, while all other peaks are removed. This process does not shuffle or exchange intensities across spectra: intensities remain attached to their original spectrum, and are only retained or removed within a replicate, and are not re-normalized. Since binned spectra are typically sparse relative to the global bin space, a replicate may remove few peaks from some spectra and many from others depending on each spectrum’s fragment composition, which motivates aggregating over many replicates.

**Figure 1.**
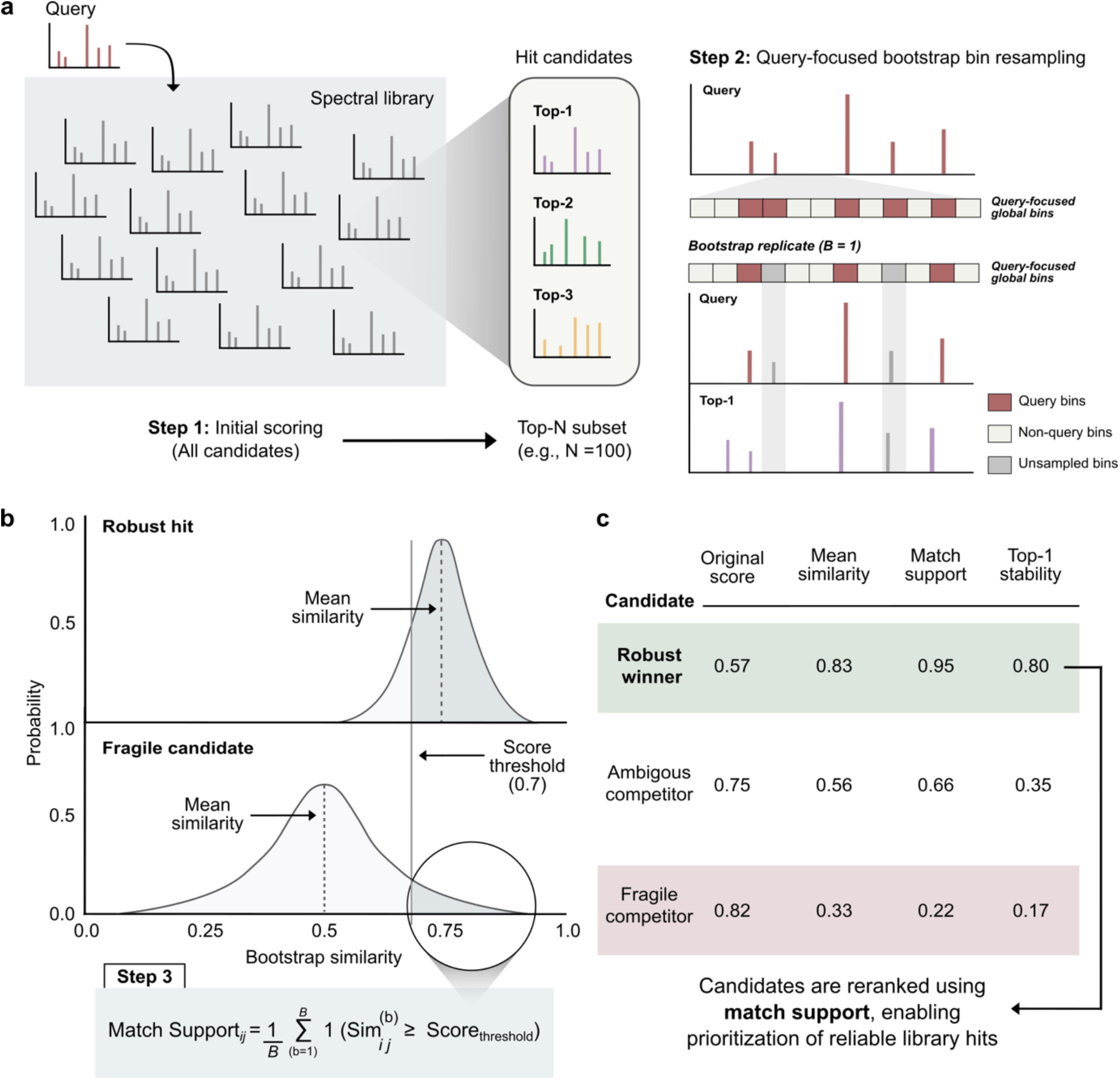
Spectral library matching with query-focused bootstrap resampling. (a) query spectra are initially scored against all library spectra, and the top-N candidates are retained. Bootstrap replicates are generated by resampling the global fragment bins derived from each query spectrum. In each replicate, query bins are resampled with replacement and spectral similarity between the query and each candidate is recomputed. (b) Repeating this over B replicates yields an empirical distribution of bootstrap similarity scores for every query–candidate pair. A reliable hit produces a narrow distribution concentrated above the score threshold, whereas a fragile candidate produces a broad, low distribution that rarely exceeds it. (c) Candidates are reranked by match support or top-k stability rather than original score, enabling prioritization of reliable library hits.

If a spectrum contains no peaks after masking, a dummy peak is inserted so that similarity computation remains well-defined; such spectra contribute effectively zero similarity for that replicate. In each replicate, the query is scored against all top-N candidates using the selected similarity metric, and the candidates are ranked. The three below-listed quantities are then aggregated across the *B* replicates for each candidate j.

1. 1.**Mean similarity**: the mean of the query–candidate spectral similarity for j across the *B* replicates, with its standard deviation reported as a measure of score reproducibility.
2. 2.**Match support**: the fraction of replicates in which the query–candidate spectral similarity for j meets or exceeds the user-defined score threshold (default 0.7) (Figure 1b). Formally, match supportⱼ = cⱼ / *B*, where cⱼ is the number of replicates in which the score for j reaches the threshold.
3. 3.**Rank stability:** the fraction of replicates in which candidate j is ranked within the top 1, top 3, or top 5, together with its mean bootstrap rank that we define as the average rank of *j* across the *B* replicates.

Finally, candidates are reranked by match support and/or rank stability rather than by their original similarity score to prioritize matches that are reproducible under perturbation of spectral evidence (Figure 1c).

### Spectral similarity metrics and configurations

Cosine and modified cosine similarities were computed using the matchms similarity implementations, with a user-defined precursor m/z tolerance where applicable. Spec2Vec similarities were computed using a model retrained for positive ionization mode spectra on curated positive mode training spectra [https://zenodo.org/records/12543129],^20^ MS2DeepScore similarities were computed using a publicly available pretrained model [https://zenodo.org/records/17826815].^21^ Model sources, versions, and download locations are reported in the Data and Code Availability section.

### Benchmarking

To systematically evaluate the effect of match support on spectral library matching, two spectral libraries were cross compared. The GNPS Natural Products Spectral Library (1,267 spectra) served as the query set, and the MSn-COCONUT dataset (15,947 spectra, see further details in the Data Availability section) served as the reference library.^17^ Ground truth relationships between query and reference spectra were defined in two steps. First, true matches were determined by comparing InChIKeys generated from the spectrum’s metadata SMILES and ignoring stereochemical information, resulting in 76 spectra. Second, we identified the number of query spectra that had an analog in the reference library by converting spectrum’s SMILES into Morgan fingerprints (RDKit, radius 2, 4,096 bits) and calculating the Tanimoto similarity between query and reference fingerprints. If this similarity was above a threshold of 0.7, two compounds were considered analogs, resulting in 142 queries with an analog but no exact match. For our analysis, only queries that had a true match or an analog in the reference database were kept, to prevent cluttering plots with a massive number of true negatives. The analysis notebook can be found at [https://zenodo.org/records/21507928].

### Perturbation experiments

We used the library_matching/library_ABUSE_GNPS_MCE_cleaned_harmonized.mgf spectra (31,742 in total) and kept only those that had an InChIKey, leaving 28,533 spectra. These were then deduplicated by grouping spectra that share the same InChIKey (first 14 characters), instrument type and adduct, keeping for each group of very similar spectra the one with the highest annotation confidence value (already defined as one of the metadata fields) and discarding the rest. This left 10,044 MS/MS spectra with 7,526 unique InChIKeys. After the filtering, each spectrum was annotated with two stratification fields for downstream analysis: (i) analogue_count, the number of other spectra in the deduplicated set whose Morgan fingerprint (radius 2, 4,096 bits, computed from SMILES) had a Tanimoto similarity ≥ 0.7 to it, spectra with a count above the median (median = 1) were treated as high-density and the rest as low-density (Figure S1); and (ii) strat_peaks, labelling spectra above or below the median peak count (median = 52) as “high” and “low” (Figure S2). The deduplicated subset was saved as library_ABUSE_GNPS_MCE_cleaned_harmonized_deduplicated.mgf. These steps are documented in the analysis notebook available at [https://zenodo.org/records/21507928].

To compare the behavior of SpecReBoot match support against the classical cosine score under controlled spectral perturbation experiments, we kept spectra retaining at least 3 peaks after removing 20 (i.e. ≥ 23 peaks). This threshold was motivated by finding a balance between having enough spectra to apply the perturbation analysis and being able to remove the highest stress level of our perturbation test. We built a perturbation grid that included four different scenarios: random peak removal (4 levels: 2, 3, 10, 20 peaks); top-k peak removal of the most intense peaks (3 levels: 1, 3, 5); addition of low-intensity peaks to simulate contaminants (3 levels: 5, 10, 20), where the added peaks intensities were drawn from the lower half of the spectrum’s intensity distribution (≤ the median) and are placed at random *m/z* between the lowest peak and the precursor; and multiplicative log-normal intensity noise (3 levels: σ = 0.3, 0.5, 0.7). Together with an unperturbed baseline and three random seeds for stochastic perturbations, this gave 34 runs in total. For each perturbed spectrum, we recorded match support and cosine score of its correct annotation relative to the baseline. The grid was run with *B* = 50, top_n = 10, score_threshold = 0.7, decimals = 2 and analog_search = True. Further details are available in run_perturbation_grid.py.

## RESULTS AND DISCUSSION

### Initial validation on a forensic DUID case

The first objective was to assess whether our query-focused bootstrapping approach could provide a per-match reliability measure that distinguishes reproducible spectral library annotations from unstable high-scoring matches. To this end, the workflow was evaluated using MS/MS data from a previously characterized forensic driving-under-the-influence-of-drugs (DUID) urine case.^14^ This dataset provided a suitable validation scenario because the sample had been extensively investigated previously, and the compounds reported in the original study could be used as reference annotations.

Previously acquired urine MS/MS spectra from the DUID case were reprocessed using the SpecReBoot library matching module and searched against the publicly available GNPS Drugs of Abuse spectral library obtained from the GNPS2 platform.5 This targeted library was selected as the initial reference space because it contains compounds directly relevant to forensic toxicology and therefore provides a controlled annotation setting with a limited number of expected true-positive compounds.

As a preliminary optimization step, different spectral preprocessing conditions were evaluated using the compound annotations reported in the original DUID study as a reference. Although the cleaning procedures implemented in matchms are optional, applying spectral cleaning to both query and reference library spectra produced the most coherent results with respect to the previously reported annotations (data not shown). Query-spectrum cleaning was therefore used to reduce the influence of low-intensity noise, while library-spectrum cleaning and harmonization were applied to minimize inconsistencies related to fragment reporting, metadata formatting, and intensity scaling across reference entries. Based on this evaluation, cleaned and harmonized query and library spectra were used for all subsequent SpecReBoot library matching analyses. After preprocessing, the urine dataset comprised 1,315 query spectra, while the Drugs of Abuse library contained 453 reference spectra.

Using the curated Drugs of Abuse library, several compounds previously reported in the DUID urine sample were recovered with high match support (0.7) and stable ranking across bootstrap replicates. These hits included cocaine, benzoylecgonine, cocaethylene, levamisole, and dextromethorphan, which combined high spectral similarity with high bootstrap-derived match support and were clearly separated from competing candidates (Figure 2a, Table S1). Importantly, the validation was restricted to compounds represented by MS/MS spectra with sufficient fragment information for bootstrap resampling; therefore, compounds associated with low-information or poor-quality MS/MS spectra were not considered suitable for reliability assessment using this approach.

**Figure 2.**
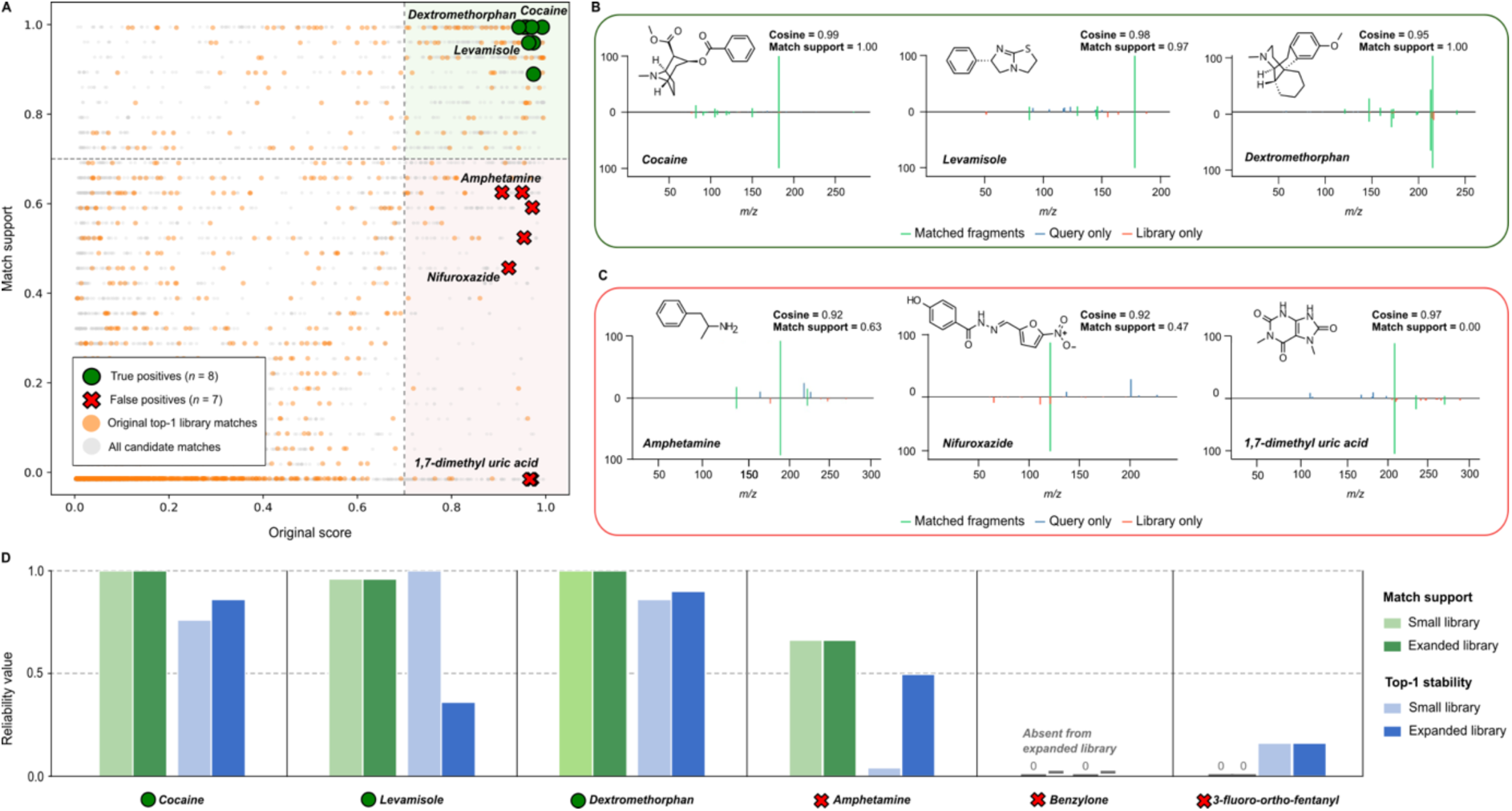
Bootstrap match support discriminates true from false annotations beyond cosine ranking. (a) Each candidate match is plotted by its original cosine score against its bootstrap-derived match support. Grey points denote all candidate matches and orange points the original top-1 library match per query. Dashed lines mark the 0.7 threshold on both axes, defining a high-reliability region (upper right, green) and a high-score/low-support region (lower right, red). Experimentally verified true positives (green circles, n = 8) cluster at high cosine score and high match support, whereas false positives (red crosses, n = 7) retain high cosine scores but collapse to low match support, separating them from genuine hits along the support axis. (b) Mirror plots and chemical structures for representative true positives, cocaine, levamisole, and dextromethorphan, and (c) representative false positives with comparably high cosine similarity but low match support, amphetamine, nifuroxazide, and 1,7-dimethyl uric acid. In each mirror plot, the query spectrum is shown on top, and the library spectrum below; matched fragments are colored green, query-only peaks blue, and library-only peaks red. (d) Comparison of match support and top-1 stability for selected compounds searched against the two evaluated reference libraries. True positives retained high match support after expansion of the search space. In contrast, top-1 stability was more sensitive to library expansion, reflecting the introduction of additional competing candidates with similar spectral evidence. Compounds absent from the expanded library search or not retained as candidate annotations are indicated near zero. Together, these results show that match support captures reproducibility of the query–candidate match, whereas top-1 stability additionally reflects competition within the candidate set.

After establishing the behavior of the method in a targeted forensic library, we next evaluated whether increasing the breadth and complexity of the reference space affected annotation reliability. The Drugs of Abuse library was therefore expanded by including a restricted GNPS-derived library and the MCE (MedChemExpress) spectral library, both available through GNPS2. Although these additional libraries are broader and less specific to drugs of abuse, they retain forensic and toxicological relevance because they include drug-related, pharmacologically active, and structurally diverse small molecules that may be absent from a dedicated forensic library.

In the expanded reference space, experimentally supported compounds still combined high cosine similarity with high match support (Table S1). Cocaine, levamisole, and dextromethorphan were consistently recovered as the top candidate with high, reproducible support (Figure 2b). In contrast, several high-scoring cosine-based matches showed low match support and unstable ranking across bootstrap replicates, indicating that their apparent similarity was not reproducible when the query spectral evidence was perturbed. This was the case for amphetamine (Figure 2c), a forensically plausible candidate that had a high cosine score (0.95) but received limited bootstrap support (0.58). Despite the initial favorable score and the forensic plausibility, the top-1 rank stability for this candidate was 0.4, meaning that in 60% of bootstrap replicates another candidate scored higher, and the apparent spectral plausibility was lost under resampling of the spectral evidence. This behavior is consistent with a match driven by a small number of shared fragments, including some of the most abundant peaks but not all, so that resampling frequently removes the evidence the match depends on while competing candidates retain enough overlap to overtake it.

The expanded reference set increased chemical coverage but also introduced a larger number of plausible competing candidates, providing a more realistic test for our approach in a complex search space. Because SpecReBoot evaluates reliability metrics within the retained top-N candidates of each query, expanding the library can change which candidates are included for the bootstrapping phase. For candidate matches retained across both searches, match support remained stable, indicating that the reproducibility of these query–candidate matches was largely preserved under resampling. In contrast, top-1 stability changed more strongly, because the expanded library introduced additional candidates that could sometimes outrank or compete with the original hit in individual bootstrap replicates. Consequently, library expansion mainly affected ranking competition among candidates, while match support remained a query–candidate specific measure of reproducibility. Across compounds, the mean absolute change in top-1 stability was accordingly ∼7-fold larger than in match support (0.22 vs 0.03; Wilcoxon signed-rank test, *p* < 10⁻³⁶; *n* = 415 compounds present in both libraries; Figure 2d).

Together, these results show that match support captures a reliability axis that spectral similarity alone does not. It penalizes high-scoring candidates whose similarity is not reproducible under resampling while preserving support for genuine annotations, even as the reference library grows and the number of competing candidates increases. In the DUID urine dataset, SpecReBoot prioritized reproducible annotations and flagged spectrally plausible but unstable candidates as less reliable matches. The expanded library analyses further showed that this behavior extends to broader forensic and metabolomic search spaces, where structurally related compounds and overlapping fragmentation patterns can confound annotation based on similarity ranking alone.

### Cross-metric match support highlights reliable endogenous metabolite annotations

We next asked whether our library matching approach could be extended from forensic drug annotations to endogenous metabolites detectable in urine. The same DUID sample was searched against the Endogenous Metabolite Spectral Library under cosine, modified cosine, Spec2Vec and MS2DeepScore spectral similarity measures. Because this library was acquired using a different instrument and chromatographic system from the query sample, this analysis provided a cross-instrument scenario in which spectral and acquisition conditions between query and reference spectra were not identical. This cross-instrument mismatch was reflected in the overall number of highly supported rank-1 annotations; among 1,303 queries scored by all four metrics, rank-1 hits with match support ≥ 0.7 amounted to 138 under cosine, 131 under modified cosine, 27 under Spec2Vec, and 70 under MS2DeepScore. These rates were lower than in the matched-instrument forensic library search where we obtained 141 rank-1 hits with match support ≥ 0.7 for modified cosine, consistent with reduced reproducibility across acquisition platforms.

Despite the cross-instrument penalty, several endogenous urinary metabolites were recovered as reliable, highly supported top-ranked annotations whose biological context further supported their biological plausibility (Table S2). For each metabolite, the same query spectrum and candidate annotation were evaluated separately under each similarity metric, allowing direct comparison of metric-specific match support values. The tryptophan–kynurenine axis showed strong but metric-dependent support: kynurenine reached high match support under cosine, modified cosine, and MS2DeepScore, with comparable cosine and modified cosine scores of 0.625 and an MS2DeepScore of 0.838. In contrast, Spec2Vec did not support the kynurenine annotation (score 0.346, support 0.00). Tryptophan showed the opposite pattern: it was fully supported under MS2DeepScore (score 0.949, support 1.00) and highly supported under Spec2Vec (score 0.712, support 0.78), but only partially supported under cosine (score 0.510, support 0.29). This pathway is activated by indoleamine 2,3-dioxygenase (IDO) during immune challenge and oxidative stress and is documented to be dysregulated by addictive substances including cocaine and opioids,^22^ with altered kynurenine metabolite concentrations observed across substance use disorders.^23^ An acylcarnitine series spanning short- to long-chain species was also recovered with high support, including butyryl- (MS2DeepScore 0.934, support 1.00), hexanoyl- (cosine 0.876, support 0.95), and palmitoyl-L-carnitine (cosine 0.991, support 1.00), consistent with urinary detection of metabolites related to mitochondrial fatty-acid β-oxidation and metabolic stress.^24^ In addition, the redox pair glutathione (modified cosine 0.951, support 0.71) and glutathione disulfide (cosine 0.867, support 0.91) were likewise recovered with high support, consistent with the role of GSH/GSSG balance as a quantitative indicator of cellular oxidative stress state.^25^

Match support also revealed instances when annotations were metric-dependent rather than a uniform consensus (Figure 3). Hypoxanthine and L-carnitine were fully supported under cosine (support 1.00 in both cases) but exhibited substantially lower support under MS2DeepScore (support 0.42 and 0.10, respectively), flagging them as annotations whose reliability depended on the scoring metric. The opposite pattern was observed for tryptophan, where cosine similarity alone resulted in a weakly supported annotation (score 0.510, support 0.29), while MS2DeepScore recovered it with full support (score 0.949, support 1.00), indicating that this embedding-based scoring model could recover an annotation that was not reliable under conventional cosine similarity. We note that match support should be interpreted within the context of the similarity metric used: it is not a fixed property of a compound, but a measure of how reproducible a query–candidate match is under a given scoring metric.

**Figure 3.**
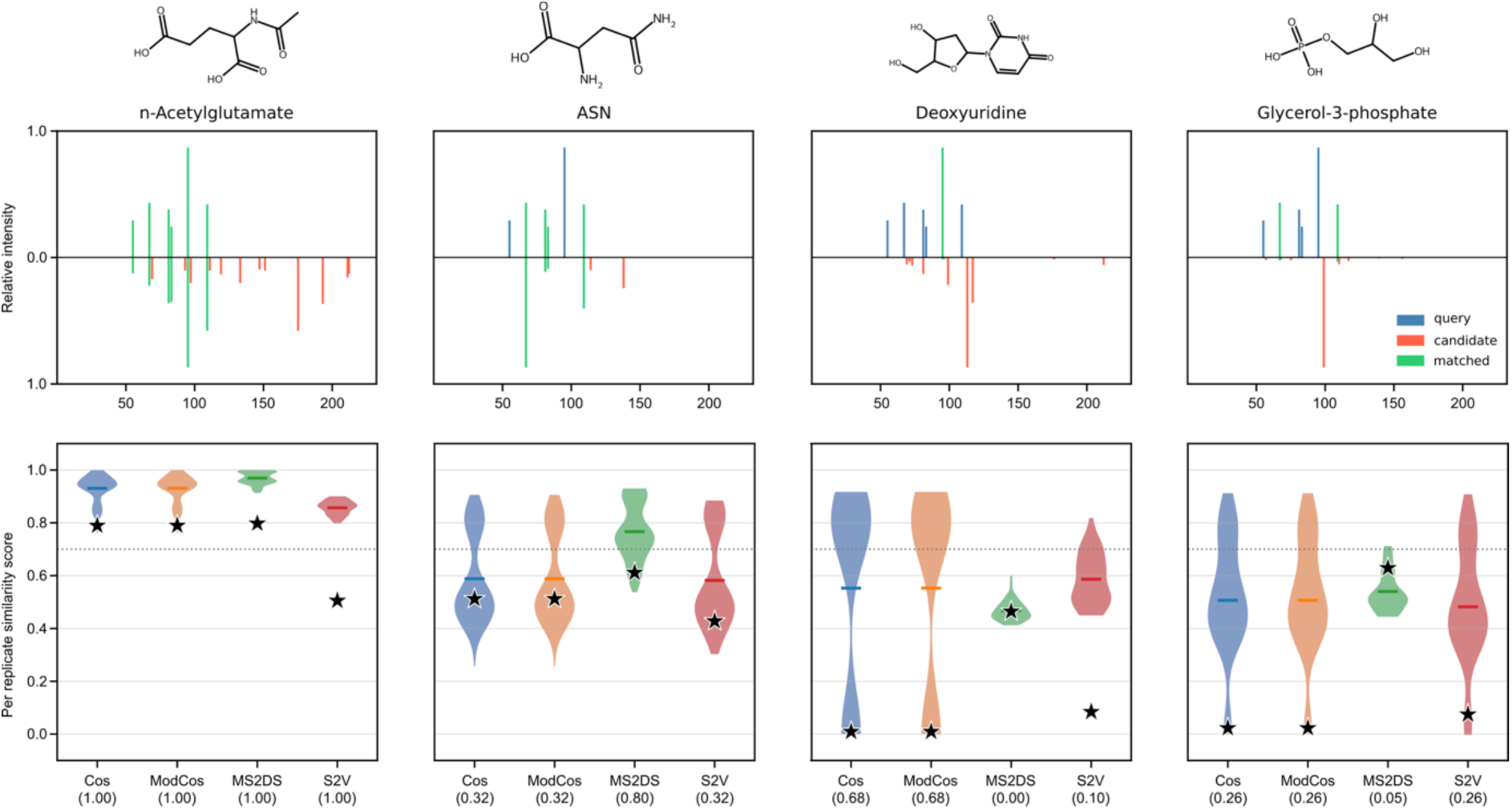
Bootstrap match support reveals cross-metric annotation behavior in a cross-instrument metabolite library search. For each of four representative query–candidate pairs, the top row shows a mirror plot of the query (above) and matched library spectrum (below), with matched fragments colored green, query-only peaks blue, and candidate-only peaks red. The bottom row shows violin plots of the per-replicate similarity score distributions across four similarity metrics, cosine (Cos), modified cosine (ModCos), MS2DeepScore (MS2DS), and Spec2Vec (S2V), derived from B = 100 bootstrap replicates. The dashed line marks the score threshold (0.7); the star (★) indicates the original non-bootstrap similarity score; values in parentheses report the match support (fraction of replicates with score ≥ 0.7) for each metric. The four examples illustrate distinct cross-metric scenarios: n-acetylglutamate (cross-metric consensus; support 1.00 across all metrics), asparagine (MS2DeepScore-specific recovery; support 0.80 under MS2DS versus 0.32 under cosine, modified cosine, and Spec2Vec), deoxyuridine (fragment-metric-specific; support 0.68 under cosine and modified cosine, collapsing to 0.00 under MS2DeepScore), and glycerol-3-phosphate (uniformly low reliability across all metrics; support ≤ 0.26).

Match support also flagged likely false positive annotations among high-scoring candidates. Since most query features in this dataset lacked independent ground-truth identities, false positives could not be systematically defined. However, precursor mass filtering served as an independent plausibility check. Among top-ranked candidates with a high raw score (cosine or modified cosine > 0.9, or MS2DeepScore > 0.75) with match support ≤ 0.2, 95% of cases showed a precursor mass differing from the query by more than 1 Da, a mass difference incompatible with correct identification regardless of the true compound identity, spanning diverse chemical classes from nucleotides (e.g. ATP) to amino acids (e.g. threonine). In one representative case, MS2DeepScore assigned vitamin B12 full support (1.00) despite a 123 Da precursor mass mismatch, while cosine and modified cosine independently converged on a different, equally low-confidence candidate, palmitoyl-L-carnitine (support 0.00, score 0.335, Δ *m/z* 155 Da). This example illustrates that neither high score nor high support alone is sufficient when precursor-mass consistency is violated; rather, match support should be interpreted together with precursor filtering, spectral metadata, and cross-metric agreement.

Cross-metric agreement therefore provided an additional axis for prioritizing endogenous metabolite spectral annotations. Metabolites supported under multiple metrics (such as kynurenine) can be prioritized as reliable candidates, whereas annotations supported under only one metric (such as hypoxanthine or tryptophan) should be flagged for manual inspection rather than accepted or rejected based on a single score. Across the 101 standard-library metabolites examined, 11% were supported by three or more metrics and 5% by only one (see Figure 3 for different examples). These results show that match support remains informative under cross-instrument mismatch, separating annotations that are reproducible across scoring strategies from those whose apparent reliability depends strongly on the similarity metric.

### Match support reduces false positive rate in library-wide benchmark

To systematically assess the effect of using match support as an orthogonal source of evidence to prioritize correct spectral library matches, 1,267 spectra from the GNPS Natural Products Spectral Library were used as queries against 15,947 spectra from the MSn-COCONUT dataset as a reference library.^17^ We restricted the analysis to queries that shared either true matches or analogs with the reference library, yielding 76 queries that had a true match and 142 with only analog matches to the reference library. Figure 4 shows the top 40 cosine matches for each of these 218 queries and the distribution of their cosine similarity and bootstrap-derived match support.

**Figure 4.**
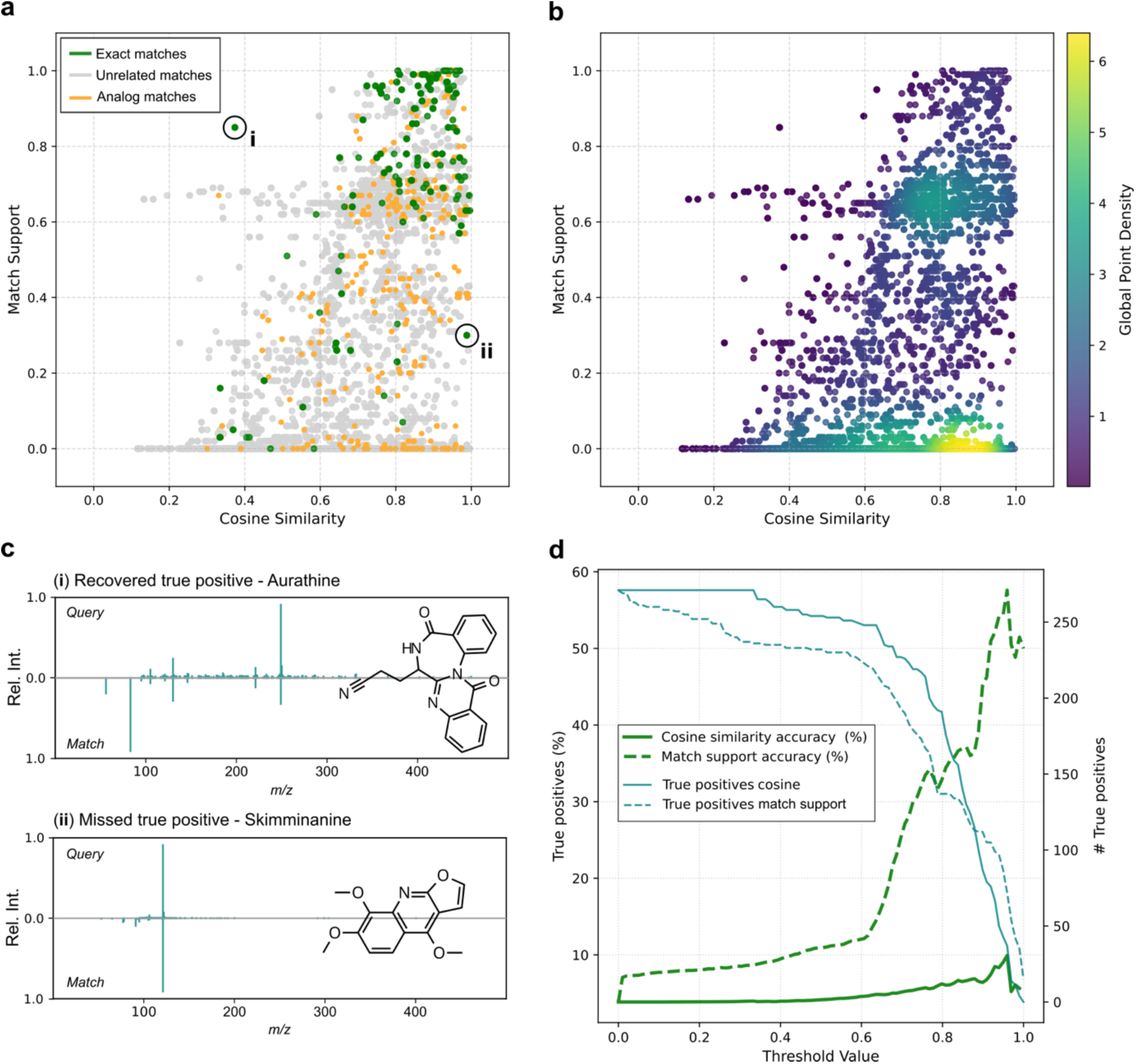
Evaluation of match support performance in cosine-based spectral library matching. (a) Top 40 cosine candidates for each of the 218 spectral queries, plotted by cosine similarity and match support. Each point represents one query–candidate match, so multiple points can originate from the same query spectrum. Matches corresponding to exact true positives are colored green, analog matches orange, and unrelated candidate matches grey. Highlighted points indicate representative true-positive outliers investigated in panel c. (b) The same set of query–candidate matches as in (a), colored by global point density on a log scale, highlighting the dominant low-support region, where many candidate matches have moderate to high cosine similarity but low support. (c) Mirror plots for the two highlighted outliers marked with i and ii in (a). Both matches are true positives, meaning query and match spectra represent the same molecule. (d). Proportion of retained matches that are true positive matches (green, left axis) and the number of true positives retained (blue, right axis), as a function of an increasing cosine or match support threshold based on the same data displayed in (a).

Notably, the cosine similarity alone was not sufficient to separate correct from incorrect spectral matches. The densest region of the scatter plot was a strip representing low-support candidates with a wide range of cosine scores, and this region contained barely any true positives (Figure 4a and 4b). The second most dense region clusters near match support ∼0.65 and included a higher number of true positives but was still dominated by false positives. Ultimately, the region with the highest density of true positives (17%) was located at the top-right corner and was characterized by a cosine similarity of ≥ 0.65 and a match support of ≥ 0.65. Notably, this analysis shows that matches with a high cosine similarity can still be false positives and that these can be further distinguished by examining their match support, considerably reducing false-positive matches. On average, exact matches moved up in rank after match support-based re-ranking (Figure S3). Under cosine-based ranking, exact matches got assigned a mean rank of 9.39 among the evaluated candidates, whereas re-ranking by match support-based improved their mean rank to 5.36. This improvement was not observed for analog matches, whose mean rank changed only slightly from 12.24 to 12.91, indicating that match support-based re-ranking preferentially promoted exact matches rather than structurally plausible alternative hits in this case.

A notable outlier in Figure 4a, representing a recovered true positive match was further investigated. This match corresponds to auranthine, a benzodiazepinone metabolite produced by the fungus *Penicillium aurantiogriseum*,^26^ which was successfully rescued by its high match support value, despite its low cosine similarity (Figure 4c). Comparison of the query and match spectrum revealed differences in fragmentation behavior, which can be linked to the fact that query and match spectra were acquired on different instruments, LC-ESI-qTof and Orbitrap, respectively. It is worth noting that this represents an isolated example, and that based on our results we did not observe the recovery of true matches with low spectral similarity as a common feature of our algorithm for cosine-based scores (Figure 4a).

Another outlier, with low match support and high cosine similarity, represented a missed true positive match, was further investigated (Figure 4c). The difference in fragmentation behavior between query and matched spectra for skimminanine, an anti-inflammatory plant metabolite,^27^ could be explained as well by the acquisition on different instruments. In this case, however, cosine similarity was high, mostly due to a shared dominant peak, which was penalized by the match support, burying this true positive match (Figure S4).

Finally, we demonstrated how prediction accuracy and the quantity of retrieved true positives were affected by varying cosine- and match support-based filtering thresholds (Figure 4d). The number of true positives retained under increasingly stringent similarity thresholds remains comparable between cosine and match support-based filtering. However, accuracy of match support-based filtering substantially outperforms cosine-based selection, showing up to 10 times more relative accuracy (threshold = 0.96). In other words, at the same recall, filtering by match support removes far more false positives than filtering by cosine.

Ultimately, these findings demonstrate match support substantially improves selectivity for cosine-based spectral library matching by reducing the dominance of any single spectral feature, filtering out false positives and in part compensating for cross-instrument fragmentation differences. The same property occasionally buries true-positive matches and demotes analogs that rely on just a few shared fragments, a trade-off that is outweighed by the substantial reduction in false positives for exact-match discovery. In practice, mass spectral similarity scores and match support are best used in tandem rather than in isolation, as spectral similarity establishes which candidates are spectrally plausible at all, and match support then indicates which of those remain reproducible when the underlying fragment evidence is resampled. Candidates that score highly on both can be prioritized with confidence, those with high similarity but low support warrant manual inspection, since they are most often driven by a single dominant fragment, and low-support candidates across the board can be deprioritized without the risk of significantly losing true annotations.

### Controlled spectral perturbation as a stress test for match support and stability rankings

Having established that match support separates reproducible from unstable annotations in the previous sections, we next assessed under which controlled spectral perturbations our query-focused bootstrapping provides the greatest benefit. For this purpose, we used the expanded reference library composed of the Drugs of Abuse library, the restricted GNPS-derived library, and the MedChemExpress (MCE) spectral library. The MCE library contains spectra from bioactive molecules, chemical scaffolds, and FDA-approved drugs, broadening the chemical search space. We retained only spectra with unique InChIKey14 identifiers, while allowing multiple adduct forms per compound, and stratified remaining entries along two axes reflecting spectrum complexity and library structural search space: peak count (median split at 52 peaks, Figure S1) and structural analog density (Tanimoto ≥ 0.7 count, median split at 1, Figure S2).

Four perturbation types were considered with the aim of reproducing distinct sources of spectral variability present in real experiments (see more details in Materials and Methods section). Following this logic, random peak removal simulated the random dropout of low-abundance fragments observed across replicate injections in DDA acquisitions.^28^ Top-k peak removal of the most intense fragments simulated the loss of diagnostic fragments caused by collision-energy shifts or precursor mass isolation issues.^29^ Added low-intensity peaks, sampled from the query’s own median-quantile intensity measurement, represent chemical noise, co-eluting contaminants, or residual fragments from previous scans.^30^ Finally, log- normal intensity noise swap (σ ≤ 0.7) simulates shifts in relative peak heights, which captures the intensity variability typically observed across instruments and collision energies. Our library matching analysis was conducted using the cosine score as the scoring metric, 50 bootstrap replicates, and allowing analog search.

Across the perturbation grid (34 conditions: 4 perturbation types × up to 4 levels × 3 seeds, each run with and without analog search), SpecReBoot’s top-1 accuracy, determined by a match support = 1.0 and the highest bootstrap-frequency candidate among precursor-mass-matched library entries, remained above 0.976 in every condition, and above 0.98 in all but the most severe perturbation case (Figure 5a). Cosine, in contrast, dropped to 0.954 under random removal of 20 peaks and collapsed to 0.869 under top-k intense peak removal at k = 5. The advantage was insignificant for intensity noise swap and for added low-intensity peaks, where both methods maintained an accuracy above 0.98. A summary per perturbation type is provided in Table S3.

**Figure 5.**
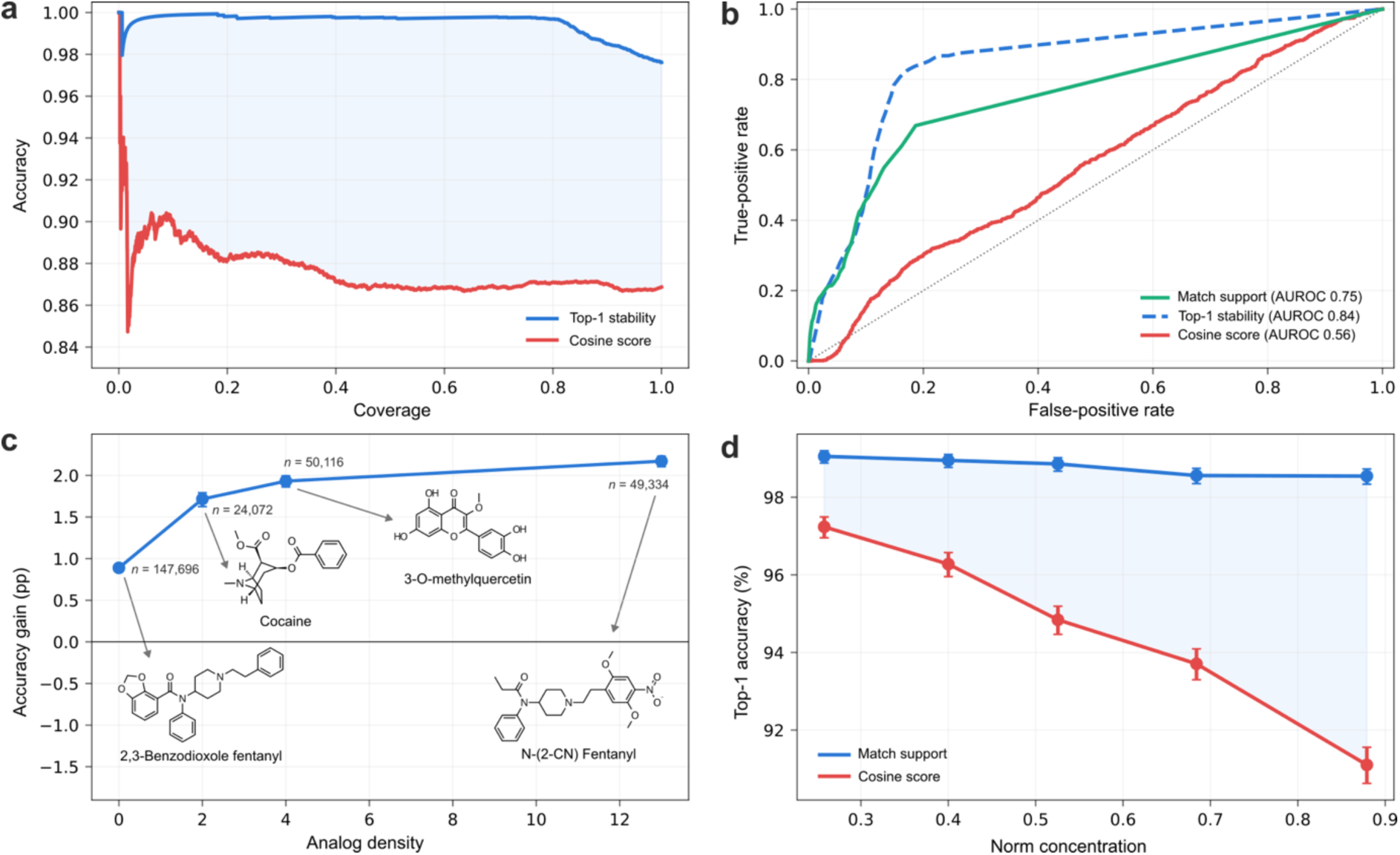
Controlled spectral perturbation of query-focused bootstrapping. (a) Risk–coverage under removal of the five most intense peaks (top-k, k = 5; *n* = 7,980). Accepting matches in decreasing order of top-1 stability holds accuracy at 0.998 over the most confident half of calls and 0.976 overall, whereas ranking by cosine score stays near 0.869. (b) Error detection across the five degradation conditions (*n* = 71,820; 867 errors, 1.21%): ROC for flagging incorrect calls, where a low reliability value should indicate an error. Match support reaches AUROC 0.75 and the cosine score of the same match only 0.56 (dotted diagonal, chance). Top-1 stability (dashed) reaches a value of 0.84, but this measure depends on the candidate space, it counts how often a candidate wins rank 1 against other competitors and can help among reliable matches to discriminate the best hit in a particular library. (c) Accuracy gain (SpecReBoot vs cosine) as a function of structural analog density (library compounds at Tanimoto ≥ 0.7), pooled over all conditions (*n* = 271,218). Points are quantile bins (median density 0, 2, 4, 13); error bars are 95% Wilson intervals. (d) Top-1 accuracy versus base-peak norm concentration across the same five conditions (*n* = 71,820). Cosine accuracy falls from 97.2% to 91.1% as concentration rises, while SpecReBoot remains near 98.5% (Spearman ρ = +1.00). Error bars are 95% Wilson intervals.

Accuracy alone, however, does not establish that the method knows when it is wrong, which is the property that matters most for spectral library matching. We therefore tested whether a low reliability value can point out an incorrect call across the perturbation conditions (*n* = 71,820; 867 errors, 1.21%). Match support, our per-match reliability metric, reached an AUROC of 0.754 for separating correct from incorrect calls, whereas the cosine score of the same match reached only 0.560, barely above random chance (Figure 5b). A high cosine similarity score therefore tells you how well two spectra matched, but not whether the match is likely to be correct – here, our introduced resampling-based scores like match support perform better. Furthermore, we note that bootstrap-derived top-1 stability resulted in a higher AUROC of 0.844, but this measure depends on the candidate space and so reflects competition among candidates rather than the intrinsic reliability of an individual match. We therefore argue that, besides looking at the reliability of spectral matches (match support), users can rely on the top-1 stability metric among reproducible matches to benefit from library space information and select the best spectral annotation.

The two stratification axes introduced at the start of this section, i.e., spectrum peak count (median split at 52 peaks), and structural analog density in the library (Tanimoto ≥ 0.7, median split at 1), illustrate these cases further (Figure S2 and S3). The advantage is smallest for spectra with a high peak count and structurally isolated from other library entries (+1.8 pp) and largest for spectra with low peak count and within dense analog neighborhoods (+8.3 pp), with the two intermediate strata falling between (+3.7 and +5.0 pp). Both axes contribute differently, analog density determines how many plausible competitors can overtake a perturbed true match (Figure 5c), while peak count determines how much evidence survives the perturbation in the first place.

This pattern is consistent with how cosine-based metrics are computed. The score is a sum over matched peaks, each weighted by the product of intensity and *m/z*, so the top few most intense peaks contribute most of the obtained score. When those peaks are removed, only a small residual overlap with the true library entry remains, and a structural analog that happens to align well with the surviving peaks can score higher. The size of this effect depends on how concentrated a spectrum’s intensity is, since cosine normalizes by each spectrum’s overall intensity magnitude, the amount the score decreases when peaks are lost is defined by the fraction of that magnitude carried by the most intense peaks. In this specific case, the library norm is highly concentrated, despite a median of 52 peaks per spectrum, the median query carries 52.5% of the norm in its base peak alone, and 93.1% in the top five peaks. Removing the top peaks therefore has the biggest effect on the score, which is precisely the top-k removal condition where cosine collapses.

Stratifying the degradation conditions by base-peak share, cosine accuracy declined consistently from 97.2% to 91.1% with increasing concentration, while SpecReBoot remained flat at ∼98.5%, widening the gap from +1.8 to +7.4 pp (Figure 5d, Spearman ρ = +1.00 across strata). This measure is largely independent of the two stratification axes above (ρ = −0.22 with peak count, ρ = +0.02 with analog density), so it captures a third and complementary property, not how many competitors exist or how many peaks remain, but how much of the score rests on the peaks a perturbation is most likely to affect. The two perturbations without significant gains are exactly those that leave the norm intact, added low-intensity peaks fall in the tail carrying ∼4% of the norm, and intensity noise rescales peaks without removing them. In both cases every candidate is re-scored on the same near-unchanged vector, the ranking is preserved, and there is no instability for bootstrap resampling to detect. Our query-focused library matching approach has therefore most added value when spectra with few peaks are searched against libraries containing structurally related candidates, a situation likely to become more common in the future with the anticipated growth of reference library sizes.^17,31,32^

## CONCLUSIONS

Reliable metabolite annotation remains a major bottleneck in untargeted metabolomics, where downstream analysis and biological interpretation are tightly linked to the candidate assignments derived, among other approaches, from mass spectral library matching. Although MS/MS spectral libraries have become essential resources for most metabolomic workflows, current spectral library matching algorithms rank candidate matches based solely on their similarity score, without a measure to judge whether each match is reliable. Here, we showed how resampling the fragment evidence of a query spectrum and recomputing its similarity across bootstrap replicates produces a per-match reliability estimate, the match support, that captures information the mass spectral similarity score does not. Across a forensic case study, an endogenous metabolite library, a natural-product benchmark, and controlled perturbation experiments, match support consistently separated reproducible annotations from high-scoring but fragile ones and identified incorrect calls where the similarity score alone was insufficient.

Two properties make our new library matching algorithm useful in practice. First, match support is a property of the individual query–candidate pair, so it remains stable as the reference library grows, whereas ranking stability (such as top-1 stability) measures shift with the candidate space. This makes the three measures complementary rather than competing: where spectral similarity establishes which candidates are spectrally plausible, match support indicates which of those survive perturbation of the underlying evidence, and rank stability then discriminates among reliable candidates within a given library. Second, the underlying statistical framework of match support is mechanistically interpretable. Its advantage is largest where a similarity score relies on a few dominant fragments, and correspondingly small where perturbation leaves those fragments intact, which allows users to anticipate when the approach will add most value, i.e., when sparse spectra are searched against libraries containing highly structurally related candidates.

Some limitations follow directly from what match support measures. As it quantifies the reproducibility of an exact match by testing for spectral overlap under resampling, it systematically down-weights analogs, which share only partial fragment evidence, and it can occasionally penalize genuine matches whose similarity depends on a single dominant peak. Match support is also metric-dependent: it reports how reproducible a match is under a given scoring function, not an intrinsic property of a compound, and agreement across metrics is therefore more informative than any single value. Finally, match support values are discriminative rather than calibrated and should be interpreted as a reproducibility measure rather than a probability of correctness.

As reference libraries continue to expand and more competing candidates appear for each query, the main question in library matching is shifting from whether a match can be found to whether it can be trusted. Our here-introduced SpecReBoot per-match and ranking-based reliability scores offer a practical route to that question by making large scale spectral library matching more transparent and its results more interpretable. As our framework is application-agnostic, we anticipate that these scores can be integrated into routine annotation workflows across metabolomics and mass spectrometry disciplines.

## DATA AND CODE AVAILABILITY STATEMENT

The SpecReBoot library matching codebase is publicly available in our GitHub repository (https://github.com/ECharria/SpecReBoot/tree/library_matching). The documentation is available in the README.md file for installation and execution. The NIH Natural Products Library and the GNPS/MCE reference libraries were obtained from the GNPS2 platform (https://external.gnps2.org/gnpslibrary), and the MSn-COCONUT dataset was obtained from.^13^ The Endogenous Metabolite Spectral Library generated for this study, together with the list of reference compounds and the associated processing notebook, is available at [https://zenodo.org/records/21394919]. The datasets used for the library-wide benchmarking and controlled perturbation experiments, as well as the analysis notebooks that include the figure creation, are available at [https://zenodo.org/records/21507928]. The urine MS/MS data used as case study were obtained from a previously published forensic toxicology investigation.^14^ Since these data are derived from a biological sample collected in a forensic context, they are subject to ethical and legal restrictions and are not publicly available, also not in anonymized form. No raw, processed, or derived files from this dataset are deposited or provided as supplementary material.

## ACKNOWLEDGMENTS

The authors thank Prof. Dr. Celia R. Berkers (Division Cell Biology, Metabolism & Cancer, Department Biomolecular Health Sciences, Faculty of Veterinary Medicine, Utrecht University) for providing access to authentic metabolite standards and for granting permission to publicly release the spectral library acquired in her laboratory. The authors thank Elena Ferri for her contribution to the initial DUID case study analysis, for the manual curation of the library-matching results in the forensic context, and for contributing to the beta-testing of the initial approach.

## SUPPORTING INFORMATION

This study is accompanied by Supporting Information file that contains Supporting Tables and Supporting Figures as referenced throughout the main text.

## AUTHOR CONTRIBUTIONS

E.C.-G. and J.J.J.vdH. conceived the concept behind the query-focused SpecReBoot library matching approach. E.C.-G. developed the methodology, implemented the computational framework, performed the data analyses, generated the figures, and led the manuscript writing. J.v.I. contributed to the codebase and beta-testing and performed benchmarking of the approach using publicly available reference spectral libraries. L.D.V. contributed the Endogenous Metabolite Spectral Library, performed the analysis of the DUID sample focused on metabolites from this library, and revised thoroughly the manuscript. L.R.T.-O. designed the controlled spectral perturbation studies, performed analysis of publicly available spectral data, and contributed to initial beta-testing of the codebase and the overall SpecReBoot project. J.J.J.vdH. supervised the project, contributed to interpretation of the results, and contributed to manuscript writing and editing. All authors reviewed and approved the final version of the manuscript.

## NOTES

J.J.J.vdH. is member of the Scientific Advisory Board of NAICONS Srl., Milano, Italy and consults for Corteva Agriscience, Indianapolis, IN, USA. All other authors declare to have no competing interests.

