## Supplementary information for "Quantifying per-match Reliability in Library Matching for Untargeted Metabolomics Workflows"

**Table S1.** Top-ranked candidate match per query reported for both reference libraries: the Drugs of Abuse library and the expanded GNPS+ MedChemExpress (MCE) library. Columns give the matched library spectrum identifier, the cosine score, the bootstrap-derived match support, and the top-1 rank stability. Δ match support and Δ top-1 stability report the change from the Drugs of Abuse to the GNPS+MCE library. Empty cells indicate that the compound was not retained as a candidate in the corresponding library (n.p.). Validation status was assigned as true positive (TP) or false positive (FP) based on the annotations reported in the original study of Ferri et al 2026.

| Compound name | Drugs of abuse (ID) | Drugs of abuse (cosine score) | Drugs of abuse (match support) | Drugs of abuse (top-1 stability) | GNPS+MCE (ID) | GNPS+MCE (cosine score) | GNPS+MCE (match support) | GNPS+MCE (top-1 stability) | Δ match support | Δ top-1 stability | Validation status |
|---|---|---|---|---|---|---|---|---|---|---|---|
| Cocaine | CCMSLIB000 11429445 | 0.996 | 1 | 0.76 | CCMSLIB0001 1429445 | 0.996 | 1 | 0.86 | 0 | 0.1 | TP |
| Dextromethorphan | CCMSLIB000 11429541 | 0.731 | 1 | 1 | CCMSLIB0000 4679246 | 0.946 | 0.98 | 0.9 | -0.02 | -0.1 | TP |
| Cocaethylene | CCMSLIB000 11429444 | 0.976 | 0.94 | 1 | CCMSLIB0001 1429444 | 0.976 | 0.94 | 1 | 0 | 0 | TP |
| Levamisole | CCMSLIB000 11429460 | 0.938 | 0.96 | 1 | CCMSLIB0001 0149293 | 0.976 | 0.86 | 0.36 | -0.1 | -0.64 | TP |
| 3-methoxymorphinan | n.p. | n.p. | n.p. | n.p. | CCMSLIB0000 4679259 | 0.973 | 1 | 1 | - | - | TP |
| Norcocaine | n.p. | n.p. | n.p. | n.p. | CCMSLIB0000 4722156 | 0.963 | 1 | 1 | - | - | TP |
| Norcocaine | n.p. | n.p. | n.p. | n.p. | CCMSLIB0000 4721848 | 0.956 | 1 | 0.92 | - | - | TP |
| Dextrorphan | n.p. | n.p. | n.p. | n.p. | CCMSLIB0000 4679248 | 0.966 | 0.98 | 1 | - | - | TP |
| Amphetamine | CCMSLIB000 09919514 | 0.76 | 0.64 | 0 | CCMSLIB0000 9919474 | 0.76 | 0.64 | 0.48 | 0 | 0.48 | FP |
| Amphetamine | CCMSLIB000 11429558 | 0.952 | 0.58 | 0.4 | CCMSLIB0000 9919474 | 0.952 | 0.58 | 0.4 | 0 | 0 | FP |
| Benzylone | CCMSLIB000 09919471 | 0.775 | 0 | 1 | n.p. | n.p. | n.p. | n.p. | - | - | FP |
| Nifuroxazide | n.p. | n.p. | n.p. | n.p. | CCMSLIB0001 0148433 | 0.923 | 0.62 | 1 | - | - | FP |
| Danegaptide | n.p. | n.p. | n.p. | n.p. | CCMSLIB0001 0145312 | 0.909 | 0.6 | 0 | - | - | FP |
| UR-144 N-(4-hydroxypentyl) | n.p. | n.p. | n.p. | n.p. | CCMSLIB0000 4721852 | 0.974 | 0.6 | 0.22 | - | - | FP |
| 1,7-dimethyluric acid | n.p. | n.p. | n.p. | n.p. | CCMSLIB0000 4679241 | 0.968 | 0 | 0.6 | - | - | FP |
| 1-methylxanthine | n.p. | n.p. | n.p. | n.p. | CCMSLIB0000 4679243 | 0.972 | 0 | 0.28 | - | - | FP |

**Table S2.** Urine sample searched against the Endogenous Metabolite Spectral Library under cosine, modified cosine, MS2DeepScore and Spec2Vec. For each metabolite the table reports the original similarity score and the bootstrap match support of its rank-1 annotation, with the number of metrics supporting it (support ≥ 0.7 spectral similarity score threshold).

| Metabolite | cosine | mod-cosine | MS2DeepScore | Spec2Vec | # metrics supported |
|---|---|---|---|---|---|
| Kynurenine | 0.625 / 0.99 | 0.625 / 0.99 | 0.833 / 1.00 | 0.329 / 0.00 | 3 |
| Tryptophan (TRP) | 0.510 / 0.29 | 0.510 / 0.29 | 0.949 / 1.00 | 0.701 / 0.74 | 2 |
| Butyryl-L-carnitine | 0.977 / 0.87 | 0.981 / 0.87 | 0.934 / 1.00 | 0.815 / 0.62 | 3 |
| Hexanoyl-L-carnitine | 0.876 / 0.95 | 0.880 / 0.95 | 0.843 / 0.63 | 0.659 / 0.89 | 3 |
| Palmitoyl-L-carnitine | 0.991 / 1.00 | 0.991 / 1.00 | 0.844 / 1.00 | 0.778 / 0.68 | 3 |
| Glutathione | 0.041 / 0.71 | 0.951 / 0.71 | – | 0.376 / 0.71 | 3 |
| Glutathione disulfide | 0.867 / 0.91 | 0.867 / 0.91 | 0.701 / 0.54 | 0.399 / 0.00 | 2 |

**Table S3.** Accuracy and error-detection performance for each of the four perturbation types, aggregated across levels and random sedes. Top-1 accuracy (%) is shown for both cosine and SpecReBoot. Δ is the SpecReBoot advantage in percentage points. The final two columns report the area under the ROC curve for flagging an incorrect SpecReBoot call from a low confidence value, using match support and top-1 stability, respectively. An AUROC of 0.5 indicates a condition with too few errors to discriminate (accuracy ~ 99.8%), so the signals gain discriminative power only where a perturbation meaningfully degrades the match, most strongly under top-k peak removal.

| Perturbation type | Level | *n* | Cosine | SpecReBoot | Δ | Match support AUROC | Top-1 stability AUROC |
|---|---|---|---|---|---|---|---|
| No perturbation | 0 | 7,980 | 99.8 | 99.8 | 0 | 0.5 | 0.5 |
| Random removal | 2–20 | 23,940 | 99.37–95.39 | ~99.7 | 0.42–4.23 | 0.71 | 0.72 |
| Top-k removal | 1–5 | 7,980 | 94.54–86.87 | 97.62 | 5.11–10.75 | 0.86 | 0.87 |
| Add intensity peaks | 5–20 | 23,940 | 99.8 | 99.8 | ~0.00 | 0.5 | 0.5 |
| Intensity noise | 0.3–0.7 | 23,940 | 99.22–98.03 | ~98.5 | ~0.15 | 0.53–0.65 | 0.79–0.84 |

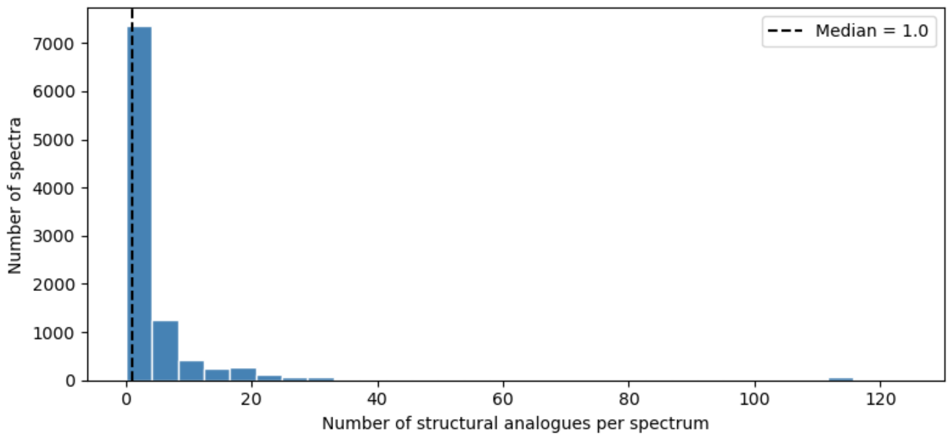

**Figure S1.** Structural analog density was defined as the number of other spectra in the dereplicated expanded reference library with a Morgan fingerprint Tanimoto similarity ≥ 0.7 to a given spectrum. The analysis was restricted to spectra with unique InChIKey14 identifiers, while allowing multiple adduct forms per compound. The dashed line indicates the median analog count of 1, which was used to divide spectra into low- and high-analog-density groups. Most spectra had few close structural analogs, whereas a smaller subset occurred in dense structural neighborhoods, representing candidates from a more competitive library search space.

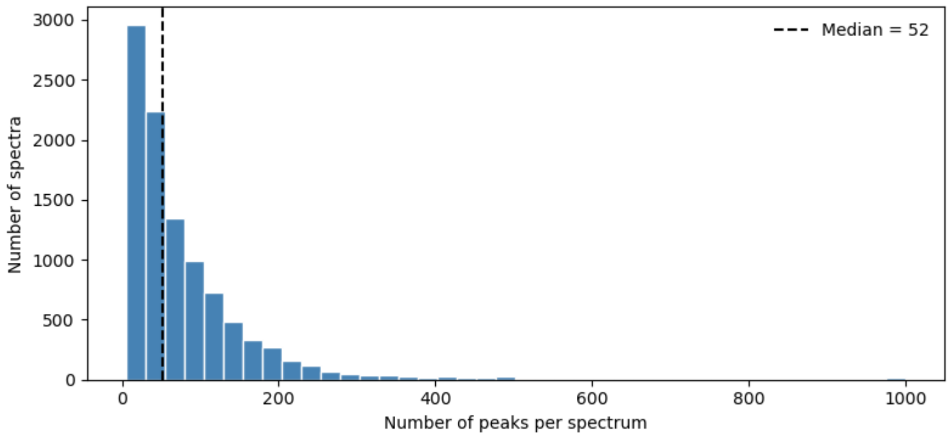

**Figure S2.** The number of fragment peaks was calculated for each spectrum retained from the dereplicated expanded reference library after filtering for unique InChIKey14 identifiers, while allowing multiple adduct forms per compound. The dashed line indicates the median peak count of 52, which was used to divide spectra into low- and high-peak-count groups. This stratification separated lower-information spectra from spectra with richer fragment evidence for evaluating the behavior of query-focused bootstrapping under controlled spectral perturbations.

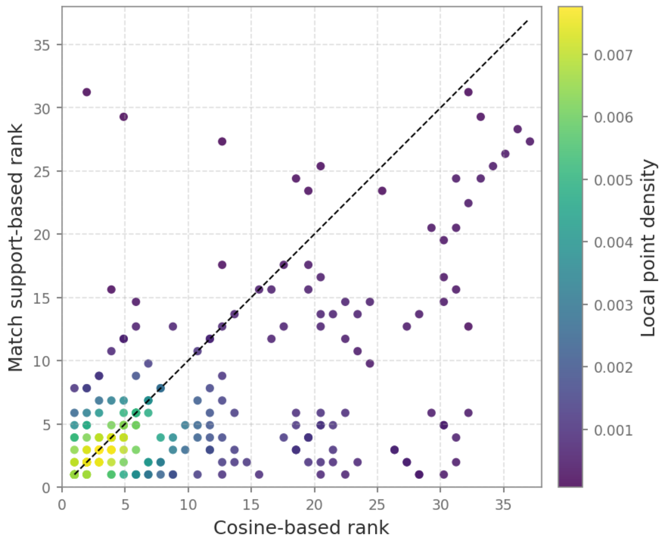

**Figure S3.** Each point represents one query spectrum for which the exact structural match was present among the evaluated candidate matches. The x-axis shows the absolute rank of the exact match under cosine-based ranking, and the y-axis shows its absolute rank after re-ranking by match support. Lower values indicate better ranks. The dashed diagonal indicates no change in rank; points below the diagonal correspond to exact matches promoted by match support-based re-ranking, whereas points above the diagonal correspond to exact matches demoted relative to cosine ranking. Point color indicates local point density, highlighting regions where multiple query spectra showed similar rank behavior. Exact matches were promoted on average, with the mean rank improving from 9.39 under cosine-based ranking to 5.36 after match support-based re-ranking.

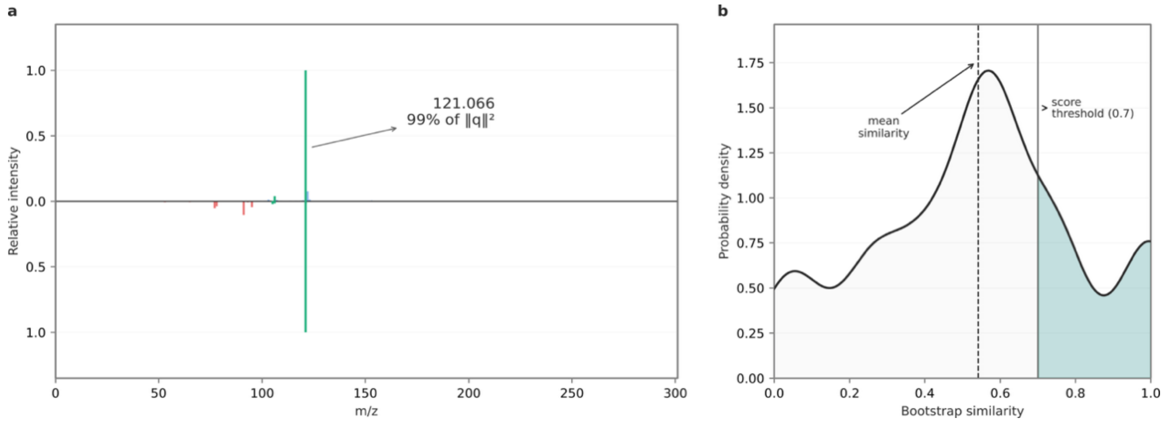

**Figure S4.** (a) Mirror plot of the query and library spectra for the skimminanine candidate match. The high original cosine similarity was driven primarily by a single dominant shared fragment at *m/z* 121.066, which accounted for 99% of the query spectral norm. Other fragments showed limited agreement between the query and library spectra, consistent with differences in fragmentation behavior likely arising from cross-instrument acquisition. (b) Distribution of bootstrap similarity scores across resampled query-fragment evidence. Although the original cosine score was high, removal or down-weighting of the dominant shared fragment across bootstrap replicates caused many replicate scores to fall below the 0.7 support threshold. This produced low match support despite a plausible true-positive annotation, illustrating that bootstrap support can penalize matches whose similarity depends on a single dominant peak rather than reproducible agreement across multiple fragments.
